# Atypical thioredoxin Patrx2 enhances alginate production in mucoid *Pseudomonas aeruginosa*

**DOI:** 10.1101/2025.07.28.667166

**Authors:** Marie M. Grandjean, Edwige B. Garcin, Moly Ba, Olivier Bornet, Christophe Bordi, Latifa Elantak, Corinne Sebban-Kreuzer

## Abstract

*Pseudomonas aeruginosa*, an opportunistic human pathogen, is known for its ability to respond and adapt to its environment, employing intricate adaptation mechanisms that can lead to the formation of complex biofilms. Redox processes play a pivotal role in bacterial adaptation mechanisms. The cytoplasm of most organisms is recognized for maintaining a reducing environment through thiol-disulfide oxidoreductases. In *Pseudomonas aeruginosa*, we have identified an unusual cytoplasmic thioredoxin named Patrx2. What sets Patrx2 apart is its active site, which contains a consensus sequence, CGHC, identical to the characteristic motif of protein disulfide isomerases (PDIs) found in eukaryotic cells. Our investigations have unveiled that Patrx2, unlike canonical thioredoxins, exhibits disulfide isomerase activity *in vitro* and displays physicochemical properties, as well as a structural conformation of its catalytic site, reminiscent of PDIs. Using a mutant transposon library, we found that the expression of *patrx2* is regulated by the alternative sigma factor AlgU, which plays a crucial role in the formation of alginate biofilms in *P. aeruginosa*. We further demonstrated strong *patrx2* expression in a mucoid strain we constructed, carrying the clinically relevant *mucA22* mutation frequently found in cystic fibrosis patients. Furthermore, our results showed a significant decrease in alginate synthesis in a *patrx2* mutant in this mucoid strain, which we attributed to its catalytic activity in a C34S variant. The study of Patrx2, an atypical thioredoxin expressed within an alginate biofilm, underscores the importance of redox regulation in adaptation mechanisms. Patrx2’s involvement in alginate biosynthesis opens new pathway for inhibiting biofilm synthesis.

**Highlights:**

- The patrx2 gene is regulated by the alternative sigma factor AlgU
- Patrx2 is strongly produced in mucoid *P. aeruginosa*
- Patrx2 is an atypical thioredoxin with disulfide isomerase activity
- Patrx2 impacts alginate production in the mucoid *P. aeruginosa* mucA22 strain, likely through redox regulation of alginate biosynthesis.

**Graphical abstract:** 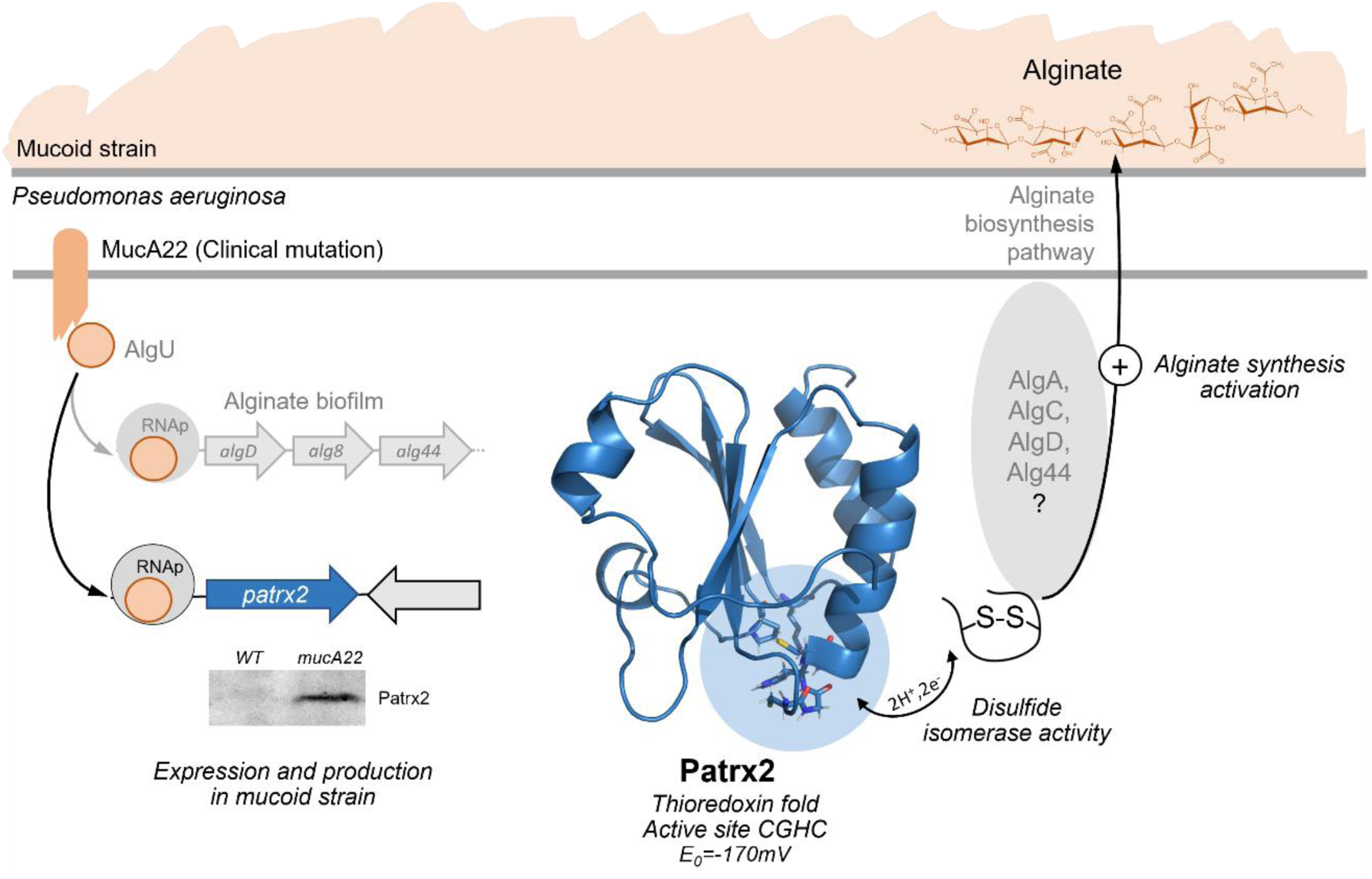

## Introduction

Disulfide bonds have been demonstrated to stabilize proteins by maintaining their overall structure through intermolecular and intra-domain covalent bonds between two cysteine residues. They are essential for the stability of secreted proteins destined for the harsh oxidizing extracellular environment and also play a role in regulating protein function. *In vivo*, the formation, reduction, and isomerization of disulfide bridges are facilitated processes [1]. Despite the vast biological diversity, most organisms on Earth use glutathione (GSH), a low molecular weight thiol compound, and thiol-disulfide oxidoreductase (TDOR) systems to perform these indispensable functions [2]. TDORs are ubiquitous in prokaryotes and eukaryotes. Cytoplasmic members of this family, such as thioredoxin (Trx) and glutaredoxin, catalyze the reduction of disulfide bonds. On the contrary, members in oxidizing cellular compartments, such as protein disulfide isomerase (PDI) in the endoplasmic reticulum, and DsbA and DsbC proteins in the bacterial periplasm, catalyze disulfide bond formation during the folding of secreted proteins [3–5].

Members of this family share a thioredoxin-fold and an active site with two conserved cysteine residues within a CXXC consensus sequence. In each TDOR subgroup, the active site has a specific conserved consensus sequence, with the XX dipeptide controlling TDOR’s redox properties and function [2,6]. Typically, the N-terminal cysteine within the active site exhibits high reactivity with a low pKa value, rendering it deprotonated at physiological pH and exposed to the solvent. Conversely, the C-terminal cysteine is buried and less reactive. The range of redox potentials varies widely, from -270mV for the most reducing to -67mV for the most oxidizing [7,8]. TDOR display a thioredoxin folding, characterized by an α/β structural motif consisting of a five-stranded β-sheet surrounded by four α helices [9]. Topological differences within the TDOR superfamily include insertions, deletions, and duplications of secondary structural elements around this motif. In TDORs like PDI, unlike prokaryotic TDORs, the protein is composed of four thioredoxin fold domains (abb’a’). Domains a and a’ exhibit catalytic activity due to the presence of the conserved CGHC motif, while domains b and b’ lack catalytic function [10]. Several residues remain highly conserved in TDOR structure, including two charged residues, an acid, and a base proximate to the active site, potentially participating in proton transfer reactions [11], as well as a cis-conforming proline situated in a loop opposite to the active site. This loop appears to influence TDORs’ redox properties and substrate binding [12–15].

In the genomes of various living organisms, you can find genes labeled as "putative thioredoxin" encoding cytoplasmic proteins with sequences characterized by "thioredoxin" motifs. These proteins often referred to as atypical thioredoxins. This nomenclature arises from their distinct characteristics: the XX dipeptide is different from that found in TDOR subgroups, and generally exhibit lower sequence identity compared to classical thioredoxins. Compiling a comprehensive list of atypical cytoplasmic thioredoxins is a challenging task, mainly due to the limited characterization of these proteins compared to their classical counterparts. Focusing on bacterial thioredoxin-like sequences (PF00085), there are currently over 24,000 annotated sequences (InterPro database). This analysis reveals the presence of 2 to 3 additional thioredoxins beyond the classical ones within each bacterium. Among them, we identify PA2694 gene encoding an atypical thioredoxin in *Pseudomonas aeruginosa*. We have designated this particular thioredoxin as Patrx2, which bears a CGHC consensus sequence, an unprecedented feature in cytoplasmic thioredoxins but commonly found in the active site of the eukaryotic PDI. The presence of this protein within *Pseudomonas* adds an intriguing layer of interest.

*Pseudomonas aeruginosa* is a Gram-negative bacterium and an opportunistic human pathogen known for its remarkable adaptability to diverse environments, both within and outside host organisms. This bacterium is a major contributor to lethal hospital-acquired infections and chronic pulmonary infections in patients with cystic fibrosis [16]. Upon invading a human host, *P. aeruginosa* can cause both acute and chronic infections [17,18]. During acute infections, *P. aeruginosa* activates genes responsible for bacterial motility and the excessive production of toxins. In contrast, chronic infections are characterized by the formation of bacterial communities enclosed within a polysaccharide-rich matrix (biofilm), providing high resistance against the host’s innate immune response and antibacterial treatments [19].

In this study, our primary objective was to characterize the structural and functional properties of Patrx2. Physicochemical characterization of Patrx2 revealed limited disulfide reductase catalytic activity but robust disulfide isomerase activity, coupled with an unusually oxidizing redox potential for a thioredoxin. We compared the structure of Patrx2, obtained using NMR spectroscopy, to canonical TDORs, noting a significant resemblance of its active site to that of PDI. Given that the presence of atypical thioredoxins likely results from an organism’s adaptation to unique environmental conditions, we investigated the genetic expression regulation of *patrx2* in *P. aeruginosa* using a transposon mutant library. This approach enabled us to demonstrate that *patrx2* expression is activated by the extracytoplasmic function sigma factor AlgU, a key regulator of alginate biofilm formation in *P. aeruginosa*. We further showed that Patrx2 is implicated in alginate production in mucoid strains of *P. aeruginosa*. Our results suggest the existence of redox regulation of alginate production in *P. aeruginosa*.

## Materials and Methods

### Bacterial strains, plasmids and growth conditions

The bacterial strains, plasmids, and oligonucleotides utilized in the study are detailed in **Table S1**. Bacteria were grown aerobically in M9 minimal media, Luria-Broth (LB) or LB agar and *Pseudomonas* isolation agar (PIA) medium used.at 37°C or 30°C. When needed, the liquid cultures were supplemented with IPTG at the final concentration of 1mM. ampicillin (Ap, 100 μg/mL), tetracycline (Tc, 15 μg/mL), gentamicin (Gm, 50µg/mL), streptomycin (Sm 2000µg/mL) and kanamycin (Km, 50 μg/mL) were added to maintain the plasmids.

### Cloning expression and purification

The cloning, expression, and purification of Patrx2 followed previously established protocols [20].

### Activity assays

The assessment of Patrx2’s activity involved insulin reduction in the presence of DTT and the refolding of scrambled RNaseA, which served as indicators of disulfide reduction and disulfide isomerase activity. These assays followed established protocols [21].

### Redox potential

To determine the redox potential, a 0.3mM protein sample was dialyzed against a 50mM potassium phosphate buffer at pH 7.0, which included 2mM of oxidized glutathione (GSSG). Patrx2 is titrated with reduced glutathione (GSH) in the range of 0–30 mM. 1H-15N HSQC spectra were recorded on a Bruker Avance III 600 MHz spectrometer with a TCI cryoprobe at 298K during the titration process, utilizing both reduced and oxidized glutathione.

### pKa determinations

pKa determinations were conducted through NMR experiments performed on a Bruker Avance III 600MHz spectrometer at 298K. NMR samples with a 1mM concentration of Patrx2, both with and without DTT, were prepared. The pH-dependent behavior of the ^13^C chemical shifts in the protein was monitored using a two-dimensional CBCACO experiment, following the methodology outlined by Bertini et al. (2006) [22]. Resonance assignments for each pH were verified using a three-dimensional HNCO experiment.

### Structure calculation of reduced Patrx2

In the process of structure calculations, 63 restraints were applied to determine backbone hydrogen bonds in the reduced Patrx2. Additionally, we incorporated 150 supplementary restraints for backbone φ and ψ dihedral angles, utilizing TALOS and the ^1^Hα, ^13^Cα, ^13^Cβ, ^13^C′, and ^15^N chemical shifts as input data [23]. Comprehensive details of the input data and structural calculation statistics are provided in Table S1. To evaluate the accuracy of the NMR models, we applied the criteria for successful structure calculation using the CYANA software [24]. Subsequently, the 20 structures with the lowest energy (total energy) were selected to create the final structural ensemble. These structures were further refined through restrained molecular dynamics using the Amber 4.1 force field within the SANDER module of Amber 10. The structure coordinates have been deposited in the Protein Data Bank under accession number 2LRC for the reduced Patrx2. The figures were generated using the PyMOL Molecular Graphics System, Version 1.2r3pre, Schrödinger, LLC.

### Construction of chromosomal variants and mutants

To generate the different chromosomal substitutions, DNA fragments corresponding to the upstream and downstream sequences (∼500 bp) of the target region we want to delete codon were amplified from PAK genomic DNA using appropriate oligonucleotide pairs **(Table S1)**. The upstream and downstream PCR products were cloned into a BamHI linearized pKNG101 suicide vector using the one step sequence and ligation-independent cloning (SLIC) strategy [25]. To perform PAK chromosome editing, the resulting suicide plasmids were introduced into the genomic DNA through conjugative transfer by a three-partner procedure using the *E. coli* pRK2013 strain. Substitution mutants were obtained by a double selection: first on PIA agar and Sm (2000 µg/ml) at 37°C, followed by NaCl-free LB agar containing 6% sucrose at 20°C. Each mutation was checked by sequencing.

### β-Galactosidase Assay

To assess β-galactosidase activity, cultures of strains harboring PAK*ppatrx2-lacZ* reporter fusions were cultivated in liquid LB medium at 37°C for overnight or 8h. Subsequently, 500 μl of each culture was centrifuged, and the resulting pellet was resuspended in 900 μl of Z buffer (comprising 10.7g/L Na2HPO4, 2H2O, 5.5g/L NaH2PO4, 0.75g/L KCl, 0.246g/L MgSO4. 7H2O, and 2.7 ml/L β-mercaptoethanol, pH 7). Permeabilization was achieved by adding 20 μl of 0.1% (w/v) SDS and 100 μl of CHCl3. Following permeabilization, a mixture of 40 μl of orthonitrophenyl-β-galactoside (ONPG) solution (4 mg/mL in Z buffer without β-mercaptoethanol) and 20 μl of permeabilized cells, diluted in 180 μl of Z buffer, was prepared. The β-galactosidase activity was then quantified and reported in Miller units (MU) [26]. All assays were conducted in triplicate, and the results were expressed as means with corresponding standard deviations.

### Transposon mutant library

The transposon mutant library was constructed following the previously established protocol [27]. Chromosomal transcription fusion PAK*ppatrx2-lacZ* was created at the *patrx2* locus as described earlier. The mini-transposon vector pBT20, employed for mutagenesis, contains a ptac promoter that can drive the expression of downstream genes depending on its orientation. Transposition of the mini-transposon was catalyzed by the Himar-1 mariner transposase. In triplicate, we spotted 50 µl of overnight cultures from the donor strain containing pBT20 (*E. coli* Sm10) and the helper strain (*E. coli* 1047/pRK2013) onto dry LB agar plates. These plates were then incubated at 37°C for 2 hours, while the recipient PAK*ppatrx2-lacZ* strain was incubated at 42°C. Subsequently, we added 100 µl of the recipient strain to each spot, and the plate was further incubated at 37°C for 4 hours. The spots were scraped off and re-suspended in 15 ml of LB medium. The resulting suspension was plated on LB agar plates containing 25 µg/mL Irgasan, 80 µg/mL X-gal, 20 µM FeSO4, and 75 µg/L Gm. A total of 3.6 × 10^4^ colonies were obtained from 180 plates, and we screened for blue colonies. The transposon insertion site was systematically determined for the mutants displaying blue colonies, while the wildtype parental clone remained white. A semi-arbitrary PCR method was employed, which involved cell lysis at 95°C, initial amplification of the sequences adjacent to the transposon insertion using a transposon-specific primer and an arbitrary primer (pBT20-1 and ARB1D-Aus primers, respectively). This was followed by a second amplification with a nested transposon-specific primer and a primer corresponding to a non-random portion of the arbitrary primer used in the first PCR (pBT20-2 and ARB2A-Aus primers, respectively) [28]. The PCR products were purified using the Macherey-Nagel NucleoSpin® Gel and PCR Clean-up kit and subsequently sequenced using the pBT20-2 transposon-specific primer.

### AlgU-binding site prediction

A position-specific scoring matrix for the AlgU sigma factor was generated using MEME (v5.5.4) from promoter sequences of known AlgU-regulated genes [29,30]. The resulting motif was formatted in MEME motif format and used as input for FIMO (Find Individual Motif Occurrences)[31]. FIMO searches were conducted on the forward and reverse strands of selected intergenic regions using default parameters, with a p-value threshold of 0.0001. Q-values were automatically computed to estimate false discovery rates.

### Colony morphologies

The Congo Red staining, enabling us to visualize exopolysaccharide production, involved depositing a 2 µL droplet from overnight cultures of various *P. aeruginosa* strains onto the agar surface, followed by overnight incubation at 37 °C. This experiment was performed on Congo Red agar plates, a solid medium consisting of Tryptone (10 g/L) and agar (1%), supplemented with Congo Red (40 mg/mL) and Brilliant Blue Coomassie (20 mg/mL).

### Carbazole assay

To assess the quantity of alginate produced by *Pseudomonas* [32], the bacterial strains were cultured in LB medium overnight at 37°C. After incubation, the culture was centrifuged at 5000 rpm for 10 minutes, and 1 mL of the supernatant was collected and placed on ice. 1 mL of cold (-20°C) 100% isopropanol was added, and the mixture was centrifuged again at 14000 rpm for 30 minutes. The resulting pellet was resuspended in 1 mL of distilled water, with occasional vortexing and sonication as needed. To create a reference alginate range spanning from 50 to 500 µg/mL, a stock solution of alginate at 0.5 mg/mL in water was used, with intermittent vortexing and sonication. For the 96-well plate alginate assay, the plate was maintained on ice to keep the temperature low. Each well was filled with 200 µL of 0.1M Borate-Sulfuric Acid, followed by the addition of 30 µL of the sample to be tested and the reference range. Subsequently, 20 µL of a 0.1% carbozole solution in ethanol was added to each well. The plate was then incubated at 55°C for 30 minutes, after which a Tecan plate reader was used to measure the absorbance of the plate at 530 nm. The alginate concentration of the test samples was calculated based on the reference range and normalized to the OD at 600 nm (OD600) of the overnight culture.

### Statistical analysis

In our study, we assessed the normality of the data using the Shapiro-Wilk test in RStudio and performed group comparisons using the Student’s t-test, assuming normal distribution (or after appropriate transformations if necessary) with a significance level of 0.05. Results are presented as means (± standard deviation), and statistical significance is reported when p < 0.05.

## RESULTS

### Patrx2 catalyses disulfide bonds formation *in vitro*

The presence of the CGHC sequence at the active site of Patrx2 similar to PDIs active sites raises questions about its enzymatic function. We have conducted a thorough examination of the redox activity of this atypical protein through *in vitro* assays.

First, we performed an insulin-reduction assay. At identical concentrations, while the canonical thioredoxin Trx1 is able to reduce insulin, Patrx2 is unable to do so (**Figure 1A**). It required an increase in the concentration of Patrx2 to determine any specific activity. However, Patrx2 showed a significantly lower activity of 0.13±0.03 ΔOD.min^-1^.mg^-1^, compared to the relative specific activity of 3.49±0.23 ΔOD.min^-1^.mg^-1^ observed for canonic Trx1 (**Figure 1B**).

**Figure 1:**
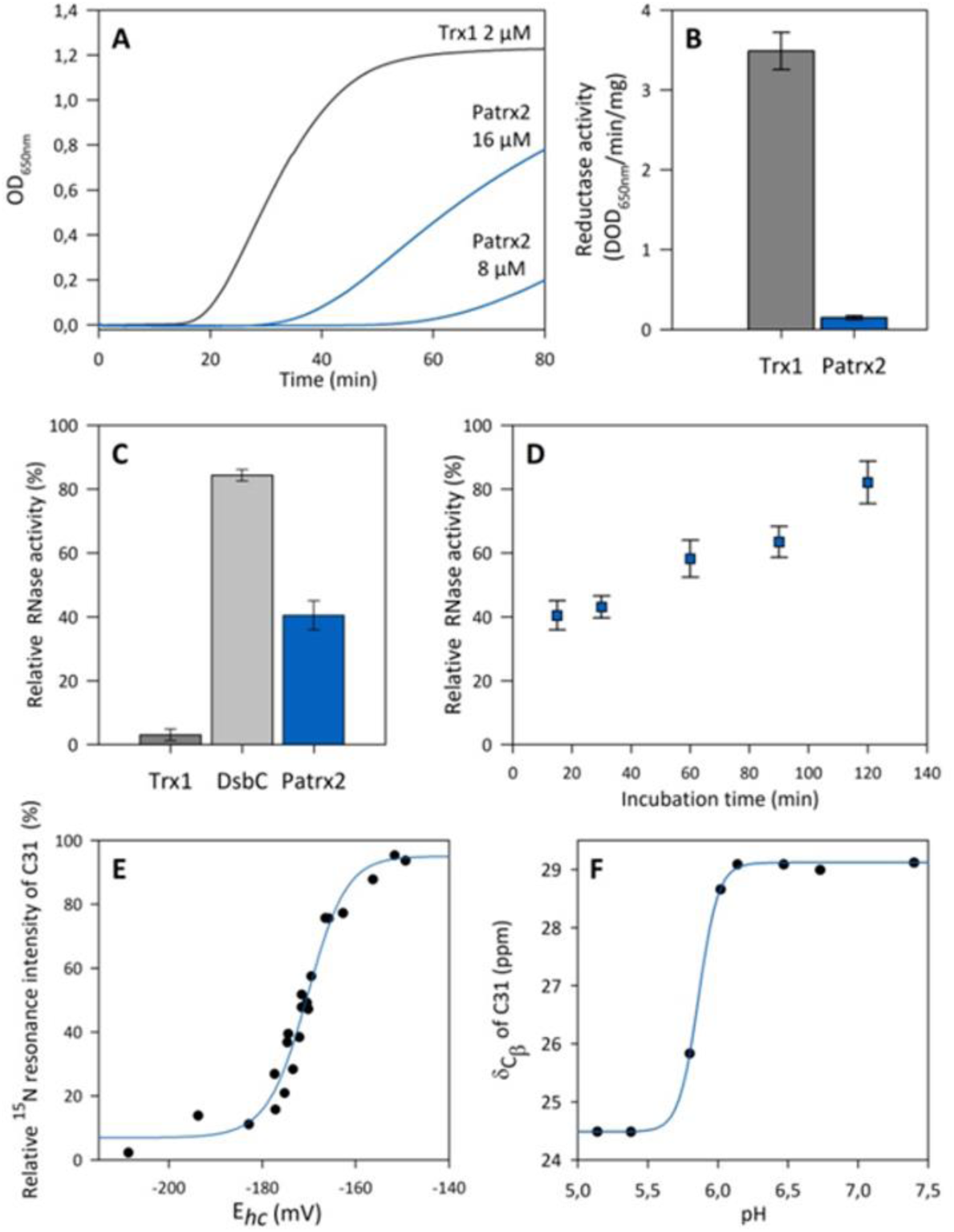
Patrx2 Structural and Physicochemical Properties. **(A-B) Disulfure reductase activity** Insulin reduction assays were conducted using Patrx2 and Trx1. Disulfide reductase activity was determined using the insulin reduction assay with different concentrations of Patrx2 (blue) or 2 µM Trx1 (grey). The catalyzed reduction of insulin (100 µM) was monitored by measuring an increase in absorbance at 650 nm, with the absorbance attributed to non-enzymatic insulin reduction by DTT (1 mM) subtracted. Specific activities were determined from at least three replicates and are presented as a histogram. The observed differences between Trx1 and Patrx2 were statistically significant (p<0.001). **(C-D) Scrambled RNaseA (ScRNase) refolding assay:** Yield of RNaseA activity of native RNaseA and reshuffling of ScRNaseA after incubation with DsbC from *E. coli,* Trx1 from *D. vulgaris* or Patrx2. For the determination of a percentage of the RNaseA activity, the mean intensity of several isolated peaks in RNA spectrum was used relative to RNA spectrum in the presence of native RNaseA. RNA spectrum in presence of ScRNaseA is used as blank. **(E) Titration of the redox potential of the disulfide bond of Patrx2**. The changes in NMR signal intensities for the NH resonances of Patrx2-C31 under oxidizing and reducing conditions in function of half- cell potential of glutathione. Black circles represent signal intensity in the oxidized state. Redox potentials were calculated using the Nernst equation from the ratio of concentrations of reduced and oxidized glutathione. Experimental data were fitted against a Nernst equation. **(F) pKa determination of C31 residue.** The pH-dependent chemical shift variation of the Cβ carbons of the nucleophilic cysteine C31 was measured and fitted to a single apparent pKa value using the Henderson-Hasselbach equation.

Subsequently, we assessed the capacity of Patrx2 to oxidise or correct incorrect disulfide bridges within scrambled RNaseA. Isomerisation will lead to RNase correct folding and therefore optimal activity. By monitoring the RNase activity through ^1^H-NMR spectroscopy, we observed the digestion of RNA during incubation. After a 15-minute incubation period, the sample containing Patrx2 exhibited a 40% increase in active RNaseA, whereas Trx1 had no impact on its activity (**Figure 1C**). Remarkably, Patrx2 demonstrated the ability to form disulfide bridges, a capability that is absent in Trx1. However, Patrx2 did not exhibit the same efficiency in refolding RNaseA as the isomerase DsbC (84%). After extending the incubation time between Patrx2 and scrambled RNaseA (**Figure 1D**), we observed an increase in RNase activity, with the enzyme reaching maximum refolding (82%) after 2h, comparable to DsbC’s refolding rate within 15 min.

### Patrx2 presents a redox potential similar to eukayotic PDIs

Next, we set out to determine the redox potential of the catalytic site of Patrx2 using NMR spectroscopy. In a previous study, the complete resonance assignment for reduced Patrx2 were performed [20]. Here, we present the backbone resonance assignment of the oxidized Patrx2 protein (**Figure S1**). The redox potential was determined through ^1^H-^15^N HSQC NMR experiments, employing the redox couple of reduced (GSH) and oxidized (GSSG) glutathione [21,33].

The intensities of the NH resonances of the oxidized Patrx2 decreased upon the addition of reduced glutathione, while the intensities of the NH resonances of the reduced form increased. The measurement of NMR signal intensities for the C31 resonance resulted in a curve (**Figure 1E**) that was successfully fitted to the Nertz equation. The redox potential for the cysteines at the active site of Patrx2 at pH 7.0 was determined to be -170.4 ± 0.5 mV.

### Patrx2 displays structural features similar to PDI active sites

Deciphering the protonation state of the residues in the active site is crucial to understand the reaction mechanism of a TDOR family protein. To investigate the pH dependency of the ^13^Cβ chemical shifts of all residues in both redox states of Patrx2, we employed a protonless NMR experiment (CBCACO). In the reduced form, the titration curves depicted in **Figure 1F** and **Figure S2** revealed pKa values of 5.8 and greater than 11.0 for the thiol groups of residues C31 and C34, respectively. Remarkably, the pKa value of the imidazole group of H33, located in the CGHC active site, exhibited an atypical behavior, shifting from 6.3 in the reduced state to less than 5 in the oxidized state of Patrx2. This alteration in the protonation state of histidine H33 could have significant implications for its activity.

To gain insight into these distinctive properties of Patrx2 and to unravel its catalytic mechanism at the atomic level, we determined the three-dimensional solution structure of this thioredoxin. The structure of reduced Patrx2 was calculated using inter-proton nOe-derived distance restraints, combined with dihedral angle and hydrogen bond restraints. We incorporated pKa values to define the protonation states of the residues during the structure refinement process. The resulting ensemble of conformers, comprising the 20 lowest-energy structures of reduced Patrx2, are depicted in **Figure 2A** (detailed structure statistics provided in **Table S2**).

**Figure 2:**
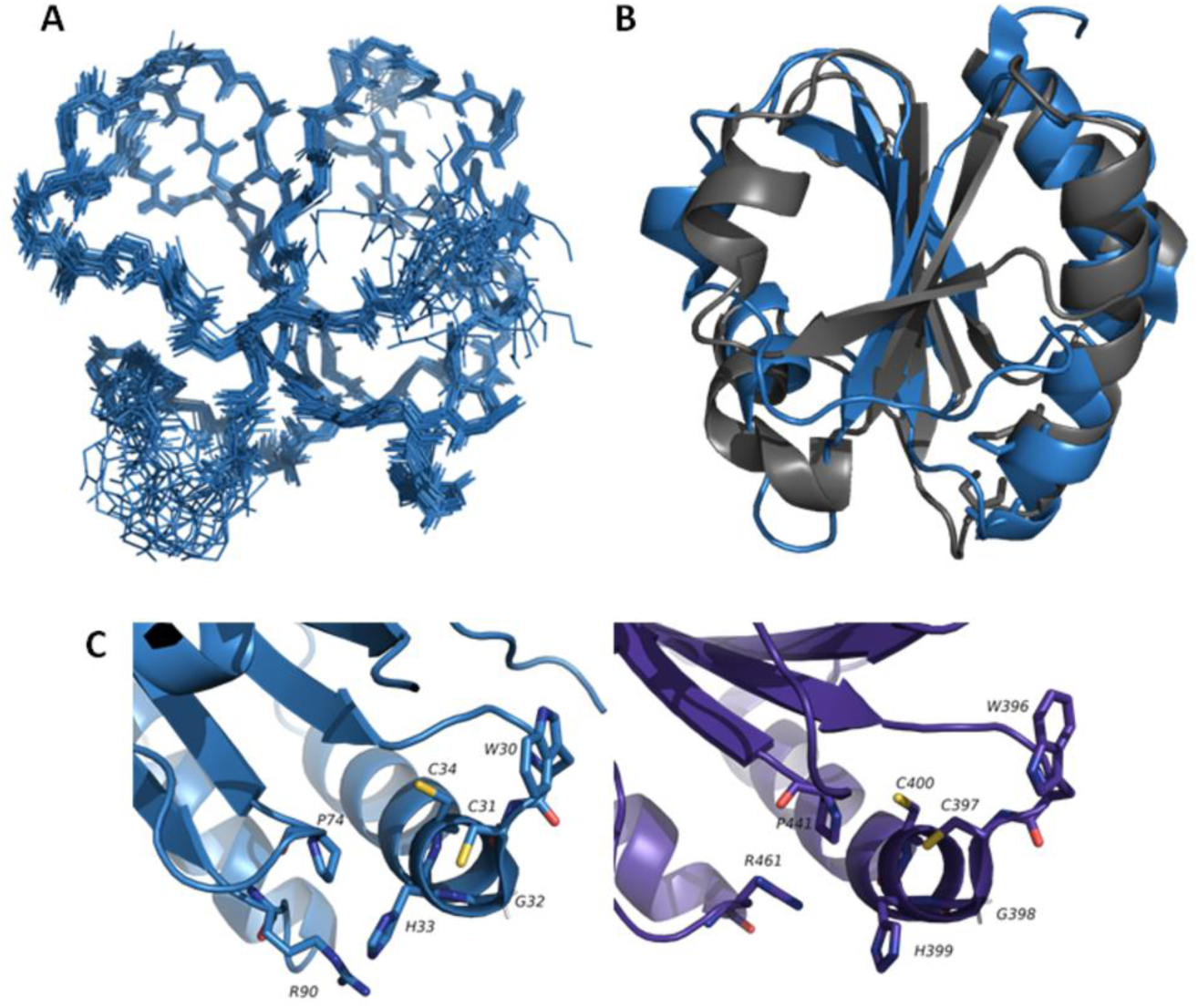
3D structure of reduced Patrx2 (2LRC)**. (A)** Superimposition of 20 representative structures of the reduced form of Patrx2 calculated with CYANA [65]. The NMR sample contained 1mM protein concentration (90% H2O, 10% D2O) in 100mM NaCl, 50mM phosphate buffer at pH 7.4 and at 290K. For reduced Patrx2, the intramolecular disulfide bond was reduced by adding DTT to a final concentration of 10mM. **(B)** Overlay of the 3D structures of Patrx2 (blue) and E. coli Trx1 (1XOA) (Grey) **(C)** Local conformation of the active site in the reduced Patrx2 (Blue) and in the human PDI (4EKZ) (purple). The active-site residues are shown and labelled. Sulphur atoms are shown in yellow, hydrogen atoms are in white, nitrogen and oxygen atoms are in blue and red, respectively.

Patrx2 adopts a characteristic thioredoxin fold [9] consisting of a five-stranded twisted central β-sheet (β1β3β2β4β5: residues 4-6 (β1), 22-27 (β2), 52-56 (β3), 75-79 (β4), and 85-89 (β5)), surrounded by four α- helices (residues 8-17 (α1), 32-47 (α2), 62-68 (α3), 94-104 (α4)). Importantly, Patrx2 exhibits a high degree of three-dimensional similarity to thioredoxins and the thioredoxin fold domain of other thiol disulfide oxidoreductases. When compared, the main chain structure de Patrx2 superimposition with that of canonical *E. coli* Trx1 reveals minimal differences (**Figure 2B**) [34,35]. However, the electrostatic surface of Patrx2 exhibits a distinct charge distribution compared to Trx1 (**Figure S3**). Specifically, we observe a slightly more positively charged surface in place of the highly conserved hydrophobic patch typically found around the Trx1 active site. Additionally, on the opposite face of the protein’s catalytic site, there is a larger negative surface area in Patrx2 compared to Trx1. This may suggest substrate specificity for Patrx2. The active site (31-CPHC- 34) of Patrx2 is located within a protruding loop between strand β2 and the N-terminus of helix α2. Specifically, the side chain of C31 is exposed at the protein surface, while the side chain of C34 is oriented toward the interior of the protein, resembling the active site configuration of reduced canonical *E. coli* Trx1. On the other hand, Patrx2 features an arginine residue (R90) in its active site identical to that of PDI. The cationic imidazole of histidine 33 is positioned between the thiolate anion of the reactive cysteine C31 and the guanidinium group of arginine R90, with mean distances of 3.0 Å and 5.28 Å, respectively.

### *patrx2* expression is regulated by the alternative sigma factor AlgU

In order to gain deeper insights into the expression of *patrx2*, we devised a global strategy. Initially, we constructed a β-galactosidase reporter strain, PAK*ppatrx2-lacZ*, in which *lacZ* was integrating at the *patrx2* locus in PAK chromosome. We found a very low β- galactosidase activity (66.8 ± 4.09 Miller Units (MU)) which suggests that *patrx2* is poorly expressed in WT strains of *P. aeruginosa* under standard growth conditions (LB media, 37°C). This observation strongly implies that *patrx2* expression is regulated, likely in response to specific environmental cues. For many years, it has been established that bacteria regulate their gene expression in response to modification in their environment [36]. This response is made through various mechanisms such as signal transduction systems involving, for example, alternative sigma factors or two-component systems. To further elucidate the regulatory mechanisms governing *patrx2* expression and identify the conditions that triggered its activation. We utilize the pBT20 system developed by Kulasekara [37], which use a mariner transposon featuring an outward promoter at one of its extremities. This transposon mutagenesis method offers the advantage of identifying genes, the disruption or overexpression of which results in increased *patrx2* expression. Our approach involved creating a transposon mutant library using the PAK*ppatrx2-lacZ* strain. This library was generated on selective media with β-galactosidase substrate (X-gal). We meticulously inspected the colonies for color variations, with a particular focus on those displaying a more intense blue colour, indicating an increase of *patrx2* locus expression. We obtained a total of 36,000 transconjugants from this mutant library, corresponding to an estimated insertion frequency of one every 174 base pairs (genome size of PAK = 6,264,404 / library size). Among these transconjugants, we specifically isolated 49 colonies that exhibited overexpression of the *ppatrx2-lacZ* gene when grown on X-gal plates. We then assessed their β-galactosidase activity in a liquid LB culture and focused on transconjugants demonstrating higher β-galactosidase activity than the PAK*ppatrx2-lacZ* strain **(Figure 3A)**. To identify the precise insertion sites of the transposons, we conducted semi arbitrary PCR using dedicated transposon primers and subsequently sequenced this PCR product to determine the insertion sites [28].

**Figure 3:**
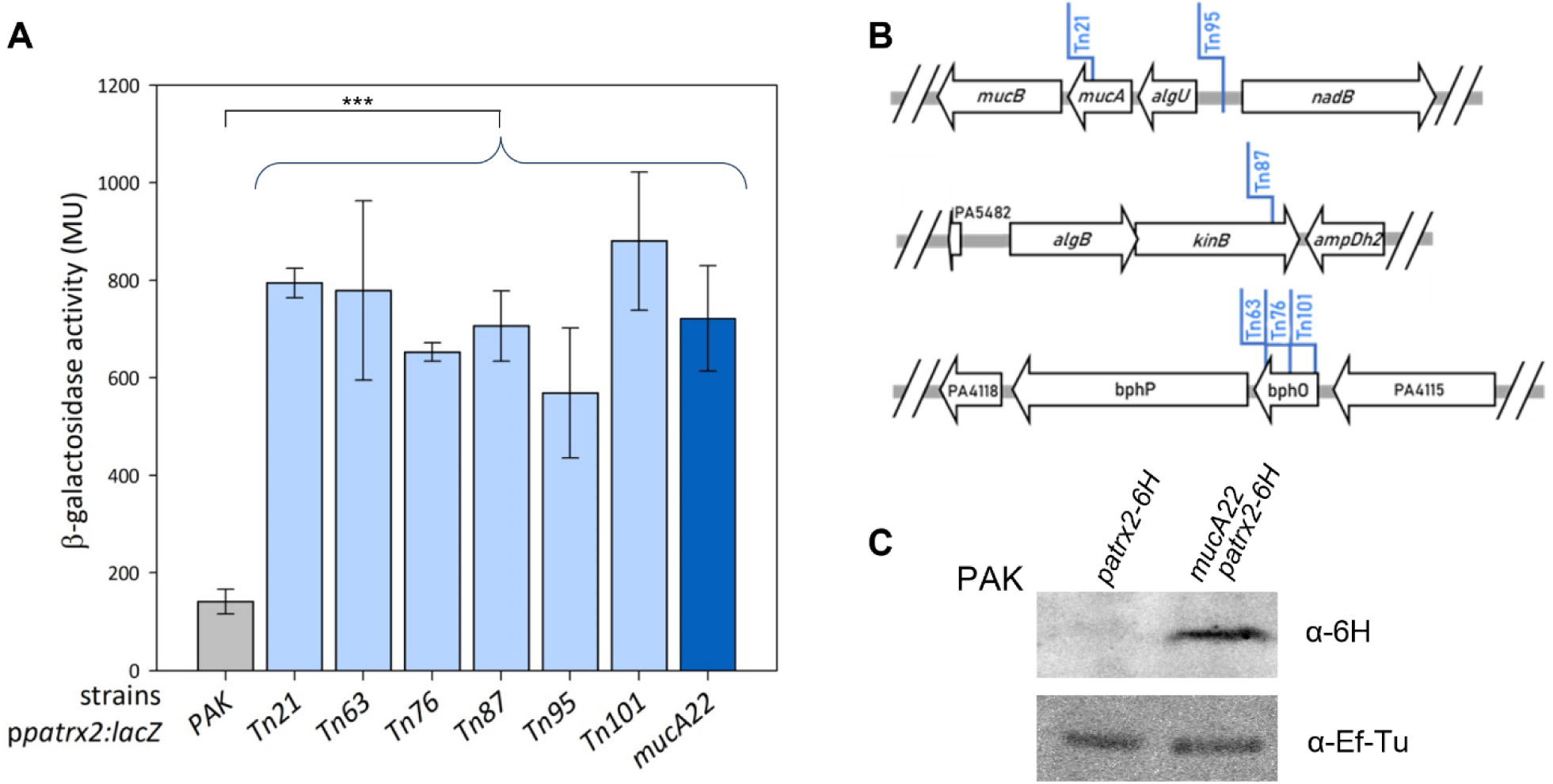
Identification of *patrx2* as part of the AlgU regulon. **(A). Expression of the *ppatrx2-lacZ* chromosomal fusion in different *P. aeruginosa* strains.** The expression of the *ppatrx2-lacZ* chromosomal transcriptional fusion was monitored in PAK, *Tn* insertion, and *mucA22* strains after 8-hour cultures in LB medium. Corresponding β-galactosidase activities are expressed in Miller units and represent the mean values obtained with standard deviations from at least 3 independent replicates. Statistical analysis of the data for all strains compared to the PAK*ppatrx2-lacZ* strain reveals statistical significance with a p-value <0.0001. **(B) Localization of transposon insertion sites in three loci of the *P. aeruginosa* PAK genome with PAO1 annotations.** The first locus (PAKAF_RS22170-PAKAF_RS22185) includes two insertions (*Tn21* to *Tn95*), the second (PAKAF_RS28950-PAKAF_RS28965) contains one (*Tn87*), and the third (PAKAF_RS04225-PAKAF_RS04240) has three (*Tn63, Tn76,* and *Tn101*). **(C) Production of Patrx2 in PAK*mucA22* by western blotting**. Patrx2 was tagged with a C-terminal 6×His tag in both PAK and PAKmucA22 strains. The strains were grown in LB medium at 37 °C with horizontal shaking at 180 rpm until the stationary phase. Cells were harvested and proteins precipitated using 20% trichloroacetic acid (TCA) at 4 °C. The pellets were washed twice with acetone, then resuspended in SDS-PAGE loading buffer, and an equivalent of 0.2 OD units was analyzed by Western blot. Proteins were detected using an anti-6×His antibody, and EF-Tu was used as a loading control.

The most frequently identified insertion locus was found upstream of the *ppatrx2-lacZ* gene and the various transposon insertion loci were presented in **Figure 3B**. Three loci have been identified. The first locus is a well-characterized operon consisting of 5 genes in *P. aeruginosa*, including two that encode the extracytoplasmic sigma factor AlgU and its anti-sigma factor MucA. In transconjugant Tn21, the transposon disrupts the coding sequence of the anti-sigma factor MucA, leading to a loss of protein function and consequently the activation of the AlgU regulon. In the second transconjugant, Tn95, the transposon is inserted in the intergenic region upstream of the *algU* operon with the transposon promoter oriented to overexpress the *algU* operon. The β- galactosidase activity of this transconjugant is lower than that measured for Tn21, but it remains significantly higher than the activity of the PAK*ppatrx2-lacZ* strain.

The second identified locus is a two-gene operon encoding the two-component system KinB (the sensor), which dephosphorylates AlgB (the regulator) [38,39]. Transconjugant Tn49 disrupts the coding sequence of *kinB* and forms mucoid colonies.

The third locus consists of an operon comprising three genes: one encoding a protein with an unknown function, the heme oxygenase BphO, and the bacteriophytochrome BphP. In this first locus, all three transconjugants obtained had the transposon inserted into the coding sequence of *bphO*, resulting in the loss of BphO function and an overexpression of *bphP*. BphP phosphorylates the transcriptional regulator AlgB, [39,40].

These findings suggest that *patrx2* expression is regulated as part of the AlgU-dependent envelope stress response. In the absence of stress, MucA sequesters AlgU at the membrane. Upon degradation of MucA, AlgU is released and activates stress-response and alginate biosynthesis genes. To test whether *patrx2* is part of this regulatory network, we constructed the mutant strain PAK*mucA22*, carrying a guanine deletion at position 425 in *mucA*, which induces a frameshift and a premature stop codon. This mutation, frequently found in clinical mucoid isolates, leads to the production of a truncated MucA protein more prone to proteolysis, thereby enabling constitutive activation of AlgU [41,42]. We introduced the *mucA* mutation into the reporter strain PAK*ppatrx2-lacZ* and measured *patrx2* expression by monitoring β-galactosidase activity, in comparison with the non-mutated strain. The results showed that β-galactosidase activity was approximately 5.3-fold higher in the PAK*mucA22ppatrx2-lacZ* strain **(Figure 3A)**. In the PAK*mucA22* background, we also tagged Patrx2 with a C-terminal 6×His tag and assessed its production by Western blot. We observed a clear overproduction of Patrx2-6His in the PAK*mucA22* strain compared to the wild-type PAK strain **(Figure 3C)**.

To investigate potential direct regulation by the sigma factor AlgU, we derived a position-specific scoring matrix based on known AlgU-regulated promoter sequences [29,30]. Using MEME and FIMO [31], we scanned 500 bp upstream of the ATG start codons of patrx2 and of the two upstream genes, pa2695 (PAK_02604) and pa2696 (PAK_02603). A putative AlgU-binding site was identified 33 bp upstream of the patrx2 start codon (p = 3.64×10⁻⁵, q = 0.0341), suggesting direct regulation by AlgU **(Figure S4)**. No significant AlgU motif was detected upstream of pa2695 or pa2696. As controls, scans of algU and algD promoter regions successfully recovered known AlgU sites with strong statistical support, validating our search strategy.

Collectively, these results confirm that Patrx2 is expressed and produced in the PAK*mucA22* strain, providing further evidence of the specific regulation of *patrx2* expression and its connection to the alternative sigma factor AlgU in *P. aeruginosa*. AlgU activates a variety of genes, including those important for biosynthesizing alginate [43]. Strains overproducing alginate are termed mucoid. The presence of atypical Patrx2 in this mucoid strain raises the question of its role and whether it could play a role in alginate biofilm formation, a hypothesis we have thoroughly investigated.

### Patrx2 is involved in alginate biofilm formation

To investigate the role of Patrx2 in alginate biofilm formation, we first constructed deletion strains of *patrx2*, complemented them onto the chromosome at the *attB* site, and generated a strain where we substituted the second cysteine of the catalytic site with serine (PAK*mucA22Δpatrx2*, PAK*mucA22Δpatrx2::attBpatrx2*, PAK*mucA22patrx2C34S*). We then assessed the five strains on Congo red agar plates to visualize exopolysaccharide production. As shown in **Figure 4**, a clear distinction is evident between the PAK strain and the mucA22 mutant strains, consistent with their mucoid phenotype. However, by direct visual inspection, no discernible differences in mucoidy among the mucA22 strains could be reliably detected using this qualitative assay. Therefore, more sensitive quantitative analyses were performed to accurately evaluate the impact of Patrx2 on alginate production. To further investigate, we subsequently quantified the amount of alginate secreted by each of these strains. For this purpose, the bacteria were grown overnight in liquid LB medium at 37°C with agitation, and the cultures were then centrifuged. Alginate content in the supernatants was determined using a carbazole assay for each sample [32]. The concentrations are reported as µg of alginate per unit of optical density at 600 nm (µg/OD600) of bacteria cultures (**Figure 4**). As expected, the non-mucoid PAK strain produced the lowest quantity of alginate, with less than 10 µg of alginate per unit of OD600, while the PAK*mucA22* strain produced a significantly higher amount of alginate, exceeding 150 µg of alginate per unit of OD600. Remarkably, the deletion of Patrx2 results in significant reduction in the amount of alginate secreted by the PAK*mucA22* strains (approximately threefold less alginate secreted per unit of OD600), and this phenotype could be complemented by inserting the *patrx2* gene into the chromosome. To further assess whether the effect of Patrx2 in biofilm formation is linked to its catalytic activity, we analyzed the *patrx2C34S* mutant. This mutant exhibits the same phenotype as the deletion mutant with the comparable loss of alginate production. These findings underscore the role of Patrx2 activity in alginate biofilm formation in *P. aeruginosa*.

**Figure 4:**
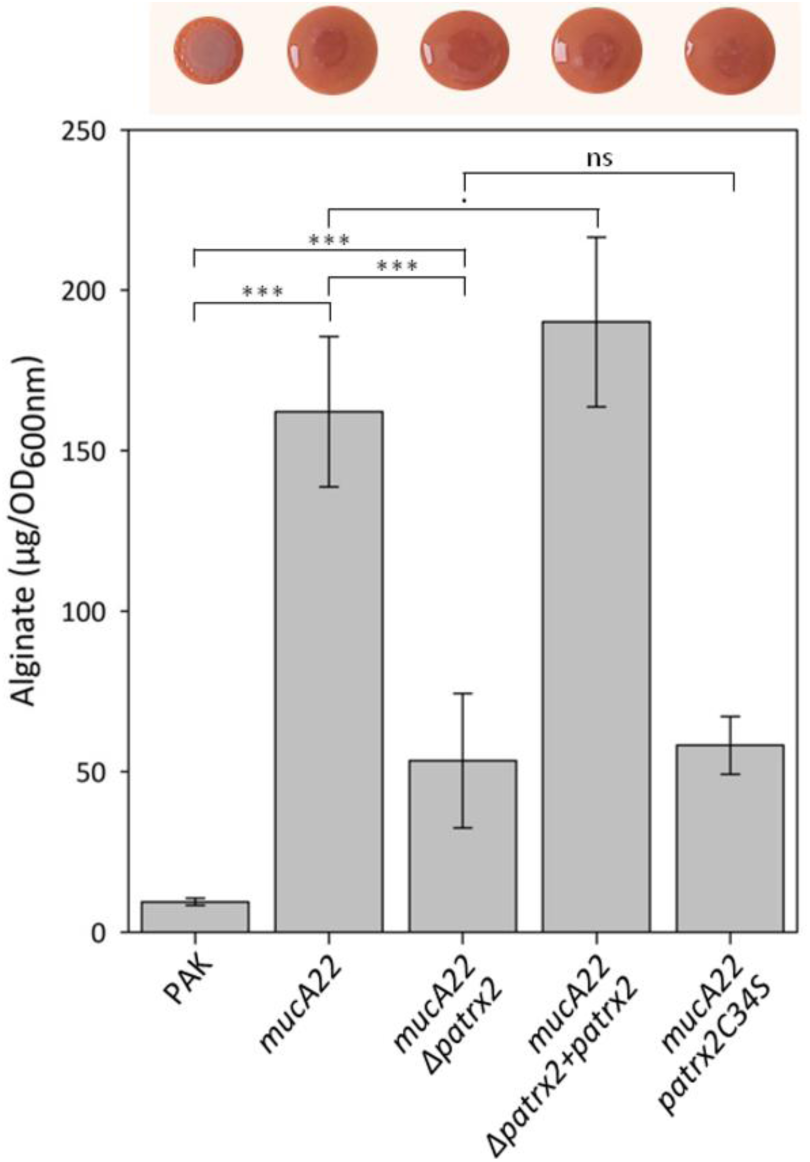
Patrx2 and alginate biofilm. **(Top):** Congo Red staining of five strains (PAK, PAK*mucA22* (*mucA22*), PAK*mucA22Δpatrx2*, PAK*mucA22Δpatrx2::attBpatrx2 (+patrx2)*, PAK*mucA22patrx2C34S*) for visualizing exopolysaccharide production on Congo Red agar plates. The solid medium consisted of Tryptone (10 g/L) and agar (1%), supplemented with Congo Red (40 mg/mL) and Brilliant Blue Coomassie (20 mg/mL). A 2 µL droplet from an overnight culture of various *P. aeruginosa* strains was spotted onto the agar surface, 24h at 37 °C. **(Bottom):** The quantity of alginate was quantified in different *P. aeruginosa* strains to assess the impact of Patrx2 on alginate production. *P. aeruginosa* strains were cultured in liquid LB over-night at 37°C. Sugars in the culture supernatants were precipitated with isopropanol, resuspended in distilled water, treated with borate to degrade complex sugars, and simple uronic acids, constituents of alginate, were quantified using carbazole, a specific colorimetric reagent for uronic acids. The absorbance at 530 nm was measured for various samples and normalized against a standard curve of known commercial alginate concentrations to obtain values in µg of alginate per OD600. The graph shows the average and standard deviation of alginate production from at least 3 independent experiments ***p<0.0001; ^•^p=0.035.

Our results indicate that the deletion of *patrx2* led to a significant reduction in the production of alginate without completely abolishing it, suggesting that Patrx2 plays a pivotal role in alginate biosynthesis. Notably, the Patrx2C34S mutant reinforces the importance of Patrx2’s catalytic activity in this process.

## DISCUSSION

*Pseudomonas aeruginosa* employs multifaceted adaptation mechanisms, presenting as single, independent free-floating cells (planktonic) or organized into microbial aggregates (biofilms). Notably, biofilm formation is a dynamic process influenced by environmental cues and serves as a protective shield against host immune responses and antibiotic treatments [44,45]. Various types of biofilms produced by *Pseudomonas* involve complex molecular networks, including critical polysaccharides such as alginate, Psl, and Pel, as well as nucleic acids and proteins, all contributing to the bacterium’s ability to survive and adapt to changing environments [46]. What role might Patrx2 play in shaping the complex adaptive strategies and virulence mechanisms of *P. aeruginosa*? To address this, our investigation into Patrx2, a cytoplasmic thioredoxin, has unveiled several atypical features, notably its distinct CGHC active site that sets it apart from classical thioredoxins. We achieved this exploration through a combination of *in vitro* assays and NMR spectroscopy, enabling us to delve into the intricate redox properties of Patrx2. This highlights that Patrx2 possesses a reductase activity approximately 26-fold lower than classical thioredoxin [47]. Patrx2 is able to both form and isomerize disulfide bridges *in vitro*, whereas DsbA never reaches a comparable level of protein refolding [48]. Patrx2 displays a redox potential of –170 ± 0.5 mV, which is more oxidizing than that of canonical thioredoxins (–270 mV) and close to that of protein disulfide isomerases (PDIs, –160 mV) [49,50]. This resemblance to PDIs is further supported by the presence of a conserved arginine residue in its active site, an amino acid characteristic of PDIs but absent in other members of the thioredoxin superfamily (**Figure 2C**). In PDIs, this arginine plays a key role in stabilizing the thiolate form of the C-terminal cysteine within the CXXC motif, which is essential for efficient catalysis [15,51,52]. Together, these findings suggest that Patrx2 is an atypical cytoplasmic thioredoxin whose structural features, particularly its redox potential and active site composition, may confer unique redox properties reminiscent of PDIs. Patrx2 is likely to have a distinct role from canonical thioredoxins in the cytoplasm, and its ability to form disulfide bonds *in vitro* could enable it to regulate specific cellular activities. It has already been demonstrated that atypical thioredoxins, like Alr2205, play highly specific roles, such as countering the toxic effects of heavy metals and H2O2 in prokaryotes [53], or as in the case of the protein BlpT, which could facilitate disulfide bond formation in a two-peptide bacteriocin, gallocin A, following secretion [54].

What regulatory networks or environmental signals might control patrx2 expression? To gain insight into the function of *patrx2*, we examined the genetic context of transposon insertions associated with its overexpression. Several of the affected loci point to established regulators of the AlgU pathway. Insertions in *bphO* (a heme oxygenase) led to *patrx2* induction. This is likely due to increased expression of *bphP*, a phytochrome sensor kinase known to stimulate AlgU-dependent gene expression in response to light and redox stress [38,39,55–57]. Insertions in *kinB* similarly led to *patrx2* overexpression. Previous studies in PAO1 have shown that *kinB* mutants induce AlgU-dependent mucoid conversion through proteolysis of MucA [40]. This effect is RpoN-dependent, and microarray data confirm that in a Δ*kinB* strain, *patrx2* expression is abolished in a Δ*kinBΔrpoN* background [41]. Finally, insertions in *mucA*, which encodes the anti-sigma factor that sequesters AlgU, are expected to derepress AlgU activity. We identified a predicted AlgU-binding site upstream of *patrx2*, suggesting direct regulation by this sigma factor. Supporting this, transcriptomic data from *P. aeruginosa* PA14 show that *patrx2* expression increases by approximately 13-fold upon AlgU overexpression [58]. The convergence of distinct regulatory pathways, *mucA*, *kinB*, and *bphO*, on *patrx2* expression, together with our confirmation of *patrx2* induction in the PAKmucA22 mutant lacking the anti-sigma factor, reinforces that *patrx2* is part of the AlgU regulon, likely responding to cell envelope stress and redox signals. This supports a potential role for Patrx2 in environmental adaptation, possibly related to biofilm formation or mucoidy. Could the expression of patrx2 in the context of this specific biofilm explain its unique redox properties? Notably, the distinct redox conditions found in biofilms are characterized by oxygen gradients. These gradients have been observed in mature biofilms, where the upper layers actively consume oxygen, thus creating distinct microenvironments within the biofilm structure [59]. Such oxygen-limited niches are particularly relevant in the context of chronic lung infections in cystic fibrosis (CF) patients, where *P. aeruginosa* often converts to a mucoid phenotype. This switch enhances bacterial persistence, in part by increasing tolerance to host-derived oxidative stress and contributing to antibiotic resistance [60]. In this context, the induction of patrx2 may reflect an adaptive response to redox fluctuations within biofilms, supporting protein folding through disulfide bond formation or isomerization in the cytoplasm. The potential link between Patrx2 and PDI-like activities may thus provide new insights into the redox adaptability and virulence of *P. aeruginosa* in hostile microenvironments such as the CF lung.

What is the potential role of Patrx2 in alginate biofilm formation? Our study represents the first step toward understanding the role of Patrx2 in alginate biofilm formation in *P. aeruginosa*, and it unveils a previously unrecognized regulatory element in this pathway. We observed a marked decrease in alginate production upon deletion of *patrx2* in the *mucA22* background. Similarly, the catalytic C34S variant of Patrx2 led to a comparable reduction, highlighting the essential role of its redox activity. Alginate synthesis involves a multiprotein complex encoded by the *alg* operon and *algC*, where most proteins act across the periplasm and envelope [61]. However, a few cytoplasmic enzymes, including AlgA, AlgC, AlgD, and the N-terminal domain of Alg44, initiate the biosynthetic process by producing and handling GDP-mannuronic acid precursors [62,63] (**Figure 5**). Interestingly, many of these cytoplasmic proteins contain cysteine residues, raising the possibility that they could be targets of Patrx2-mediated thiol-disulfide exchange or isomerization. Supporting this hypothesis, previous work showed that ebselen (Eb), a cysteine-targeting compound, inhibits alginate production downstream of transcription, presumably by acting on Alg44 or other cysteine-containing proteins [64]. Given that Patrx2 shares a thioredoxin fold and is essential for full alginate production, we propose two non-exclusive hypotheses: either Patrx2 and Eb target overlapping protein substrates in the cytoplasm, or Eb may directly inhibit Patrx2 itself—much like the functional loss observed in the C34S variant. Future work will be required to determine whether Patrx2 directly interacts with cytoplasmic enzymes of the alginate biosynthetic pathway, or if it functions more broadly as a redox sensor integrating oxidative stress signals with biofilm matrix production.

**Figure 5:**
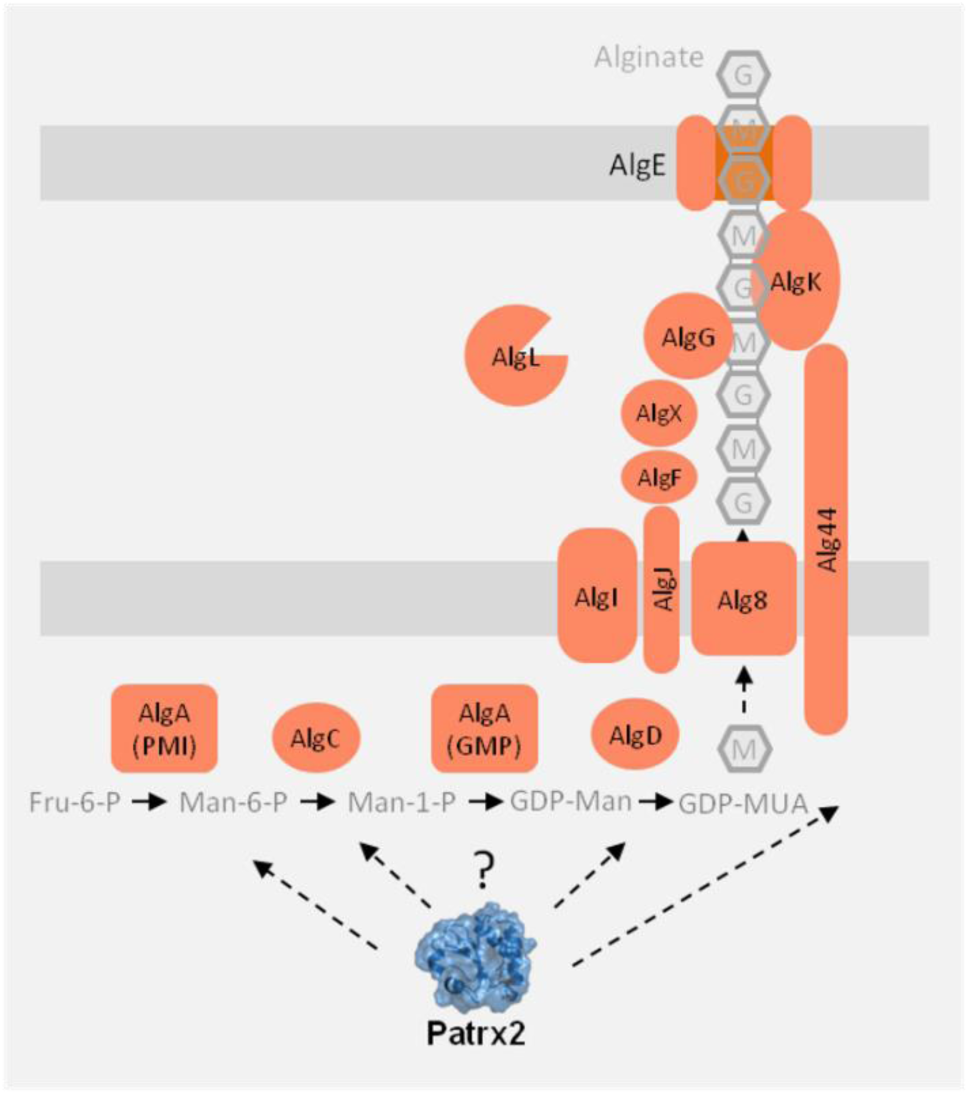
Hypothetical Role of Patrx2 in the Alginate Synthesis Pathway.

In *Pseudomonas aeruginosa*, the alginate synthesis pathway is encoded by the *algD-algA* operon and the *algC* gene [61]. The proteins are localized within the cell, including the cytoplasm, inner membrane, periplasm, and outer membrane. Soluble cytoplasmic proteins AlgA, AlgC, and AlgD provide the activated nucleotide sugar precursor, GDP-mannuronic acid, while other proteins constitute an envelope-spanning multiprotein complex. Alg8 and Alg44 play essential roles in alginate polymerization [62,63]. O-acetylation in the periplasm is catalyzed independently by AlgJ and AlgX [66–68], with AlgI and AlgF providing acetyl groups [61]. AlgG, AlgX, and AlgK assist in guiding alginate through the periplasm for secretion via AlgE [68,69].

In summary, our results demonstrate that Patrx2 is an atypical cytoplasmic thioredoxin with disulfide isomerase activity *in vitro*. It is strongly induced under the control of the stress-response sigma factor AlgU, particularly in mucoid strains, such as those frequently isolated from chronic infections, highlighting its potential clinical relevance. Patrx2 plays a key role in alginate production in mucoid P. aeruginosa strains. These findings identify Patrx2 as a novel regulatory component of the alginate biosynthetic machinery, likely acting through redox control of specific cytoplasmic targets. Future work should focus on identifying these molecular targets, which may include enzymes directly involved in alginate synthesis. Beyond its mechanistic interest, understanding the function of Patrx2 may open new avenues for targeting biofilm formation and persistence in *P. aeruginosa* and related pathogens.

## Acknowledgments

We would like to express our gratitude to Dr. Françoise Guerlesquin for her participation and support throughout, as well as to Prof. James Sturgis and the team of Prof. Sophie Bleves for their insightful discussions. The language and readability of this document were enhanced with the assistance of Generative AI (Chat-GPT 3.5).

## Funding

This work received support from the french government under the France 2030 investment plan, as part of the Initiative d’Excellence d’Aix-Marseille Université – A*MIDEX AMX-22-RE-AB-006

## Abbreviations

PDI: protein disulfide isomerase
Trx: thioredoxin
TDOR: thiol-disulfide oxidoreductase
GSH: reduced glutathione
GSSG: oxidized glutathione
X-gal: 5-bromo-4-chloro-3-indolyl-beta-D-galactopyranoside
OD600: optical density at 600nm

## Supplementary Data

**Figures S1:**
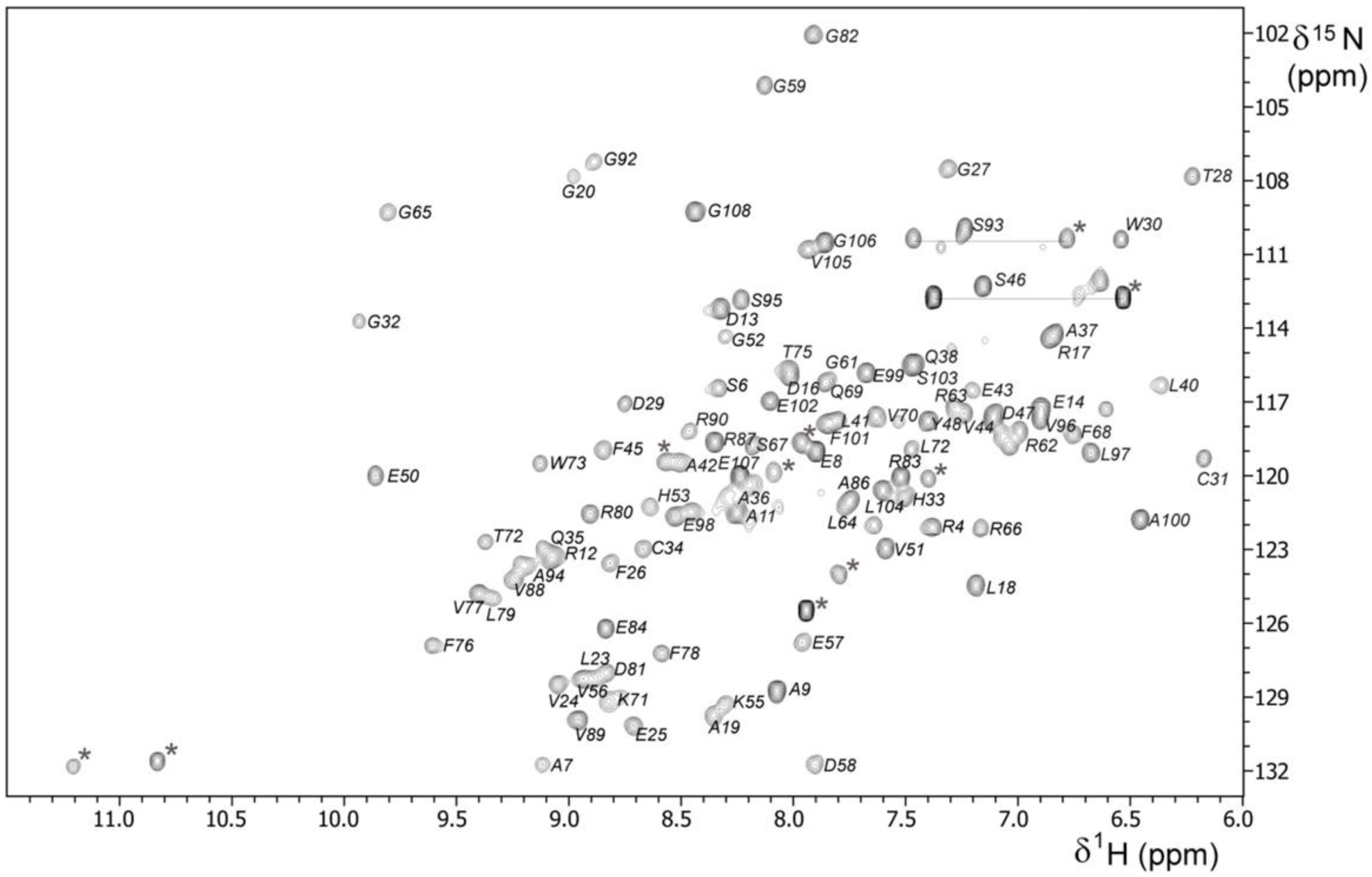
^1^H,^15^N-HSQC spectrum of oxidized Patrx2 in 150 mM NaCl, 10 mM KPO4 buffer pH8, 10 % D2O, at 300 K on a Bruker Avance III 600 MHz spectrometer. The backbone ^1^H,^15^N correlations are labeled according to the sequence. Side chain amine resonances are indicated with grey labels for Gln and Trp residues, and with grey star for Arg residues. Side chain resonances of Gln residues are connected by horizontal lines

**Figure S2:**
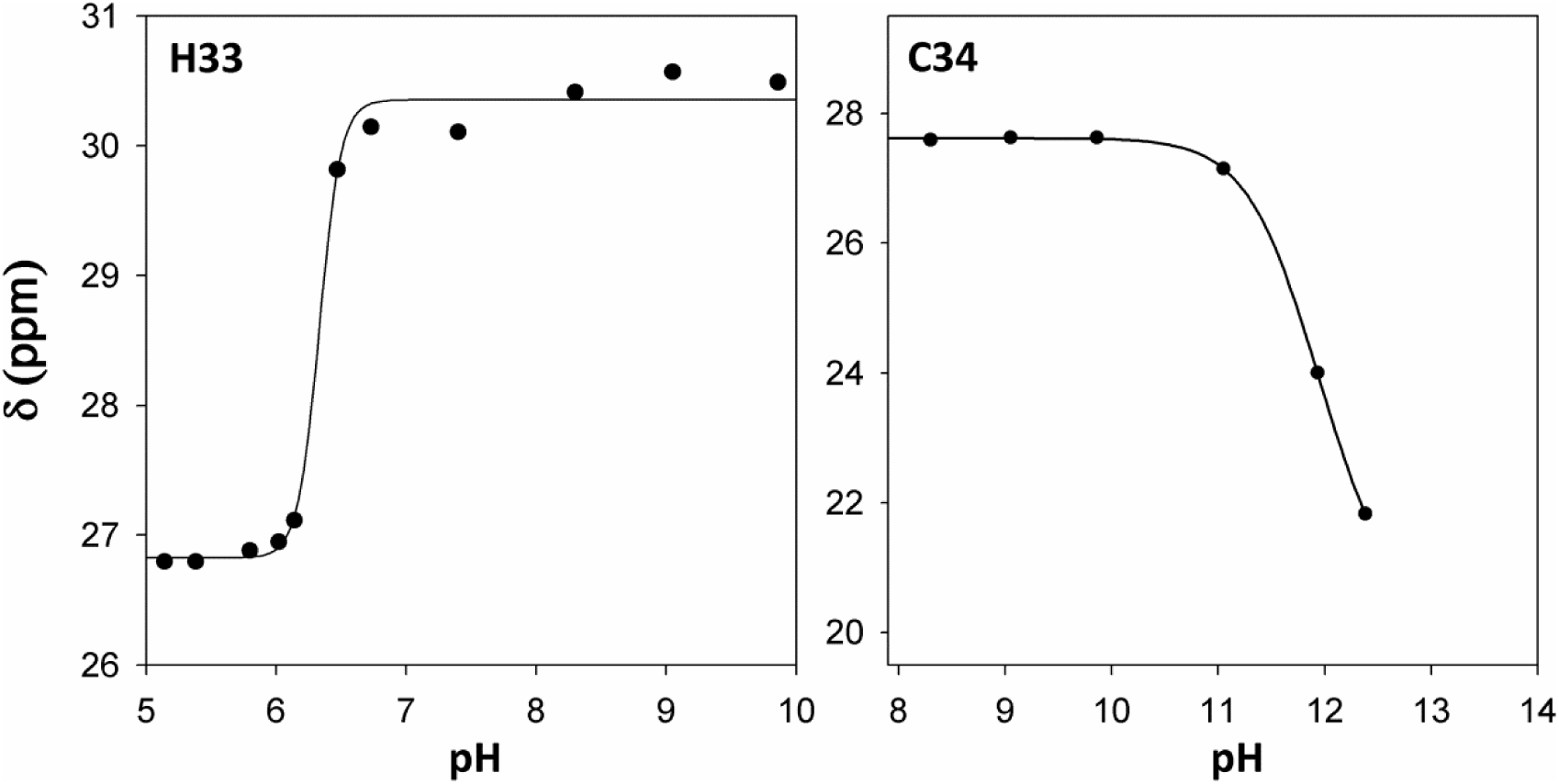
pKa determination of Patrx2 ionisable residues. ^13^C NMR chemical shift pH titration curves for several Cβ resonances. pKa determination of the histidine H33, and cysteine C34 in the reduced form of Patrx2. The pH dependent chemical shift variation of the Cβ carbons was measured and fitted to one apparent pKa value using the Henderson-Hasselbach equation.

**Figure S3:**
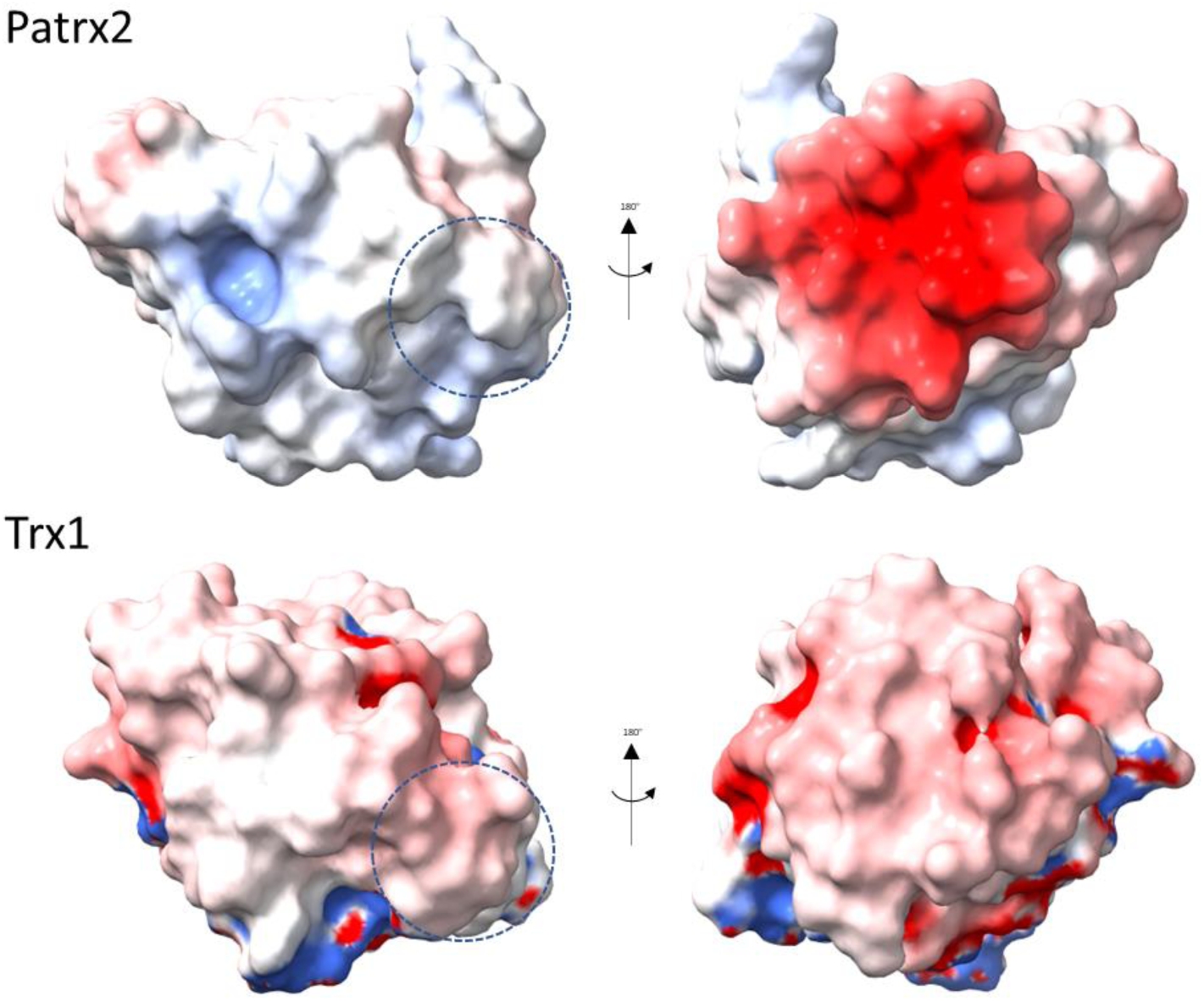
Electrostatic surface potential comparison between Patrx2 and canonical E. coli Trx1. Electrostatic potential maps were generated using APBS after structure preparation with PDB2PQR, and visualized in ChimeraX using the command color electrostatic. Surface potentials are shown from two opposite orientations (180° rotation around the vertical axis) for each protein. Red and blue correspond to regions of negative and positive electrostatic potential, respectively, with the scale ranging from –10 kT/e (red) to +10 kT/e (blue). Compared to Trx1, Patrx2 exhibits a markedly different charge distribution: a more positively charged surface near the active site region (left panels, dashed circle), and an extended negatively charged region on the opposite face (right panels). These differences may reflect functional divergence between the two thioredoxins. Electrostatic surface potential representations of Patrx2 and E. coli Trx1 in two similar orientations; the CXXC site is encircled by a dashed circle for both proteins.

**Figure S4:**
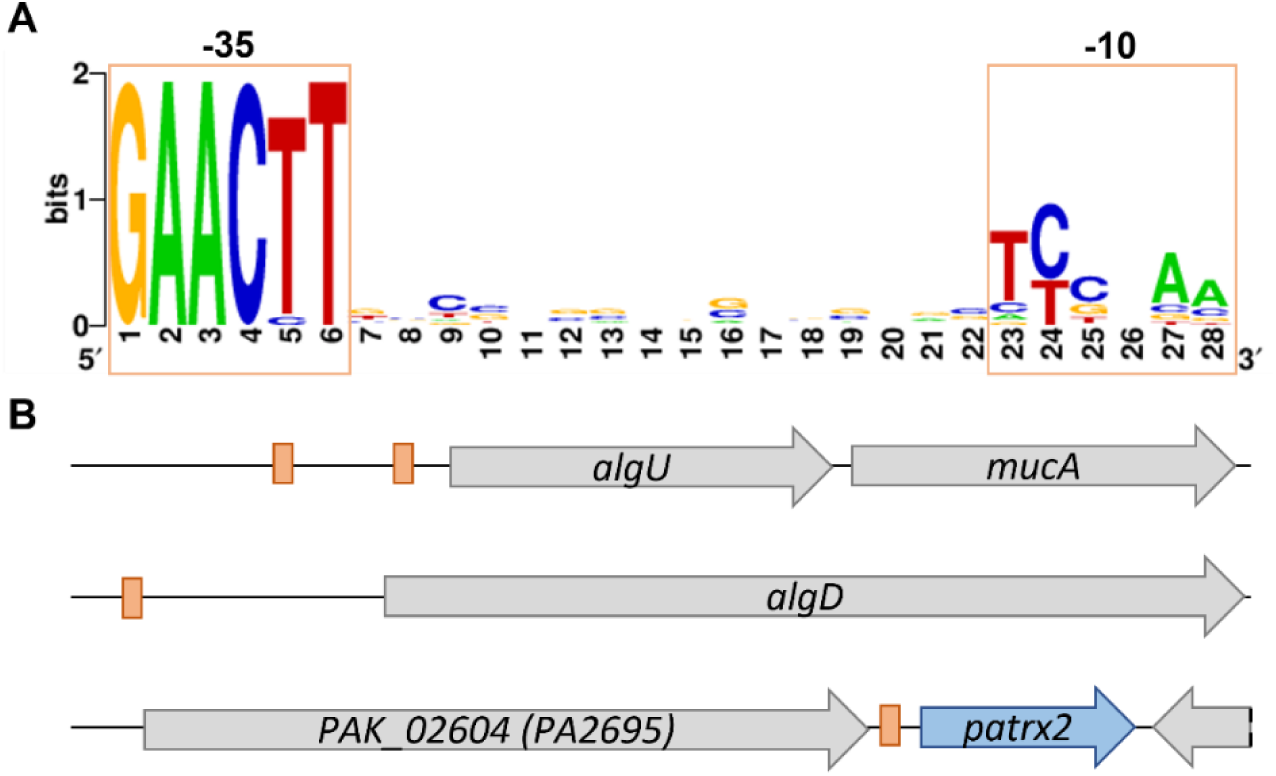
Identification of a putative AlgU binding site upstream of *patrx2*. **A.** Sequence logo of the AlgU binding motif defined using MEME based on published AlgU-dependent promoter. Conserved -35 (left box) and -10 (right box) elements are highlighted in green. **B.** Genomic context of known AlgU-regulated genes (*algU*, *algD*) and the *patrx2* locus in *P. aeruginosa* PAK. Green boxes indicate predicted AlgU-binding sites identified using FIMO. For *algU*, binding sites were detected 58 bp (p = 2.02×10⁻⁵, q = 0.0252) and 242 bp (p = 2.67×10⁻⁷, q = 0.001) upstream of the start codon. The *algD* site is located 372 bp upstream (p = 1.74×10⁻⁵, q = 0.0252), and the *patrx2* site lies 33 bp upstream (p = 3.64×10⁻⁵, q = 0.0341).

**Table S1:**
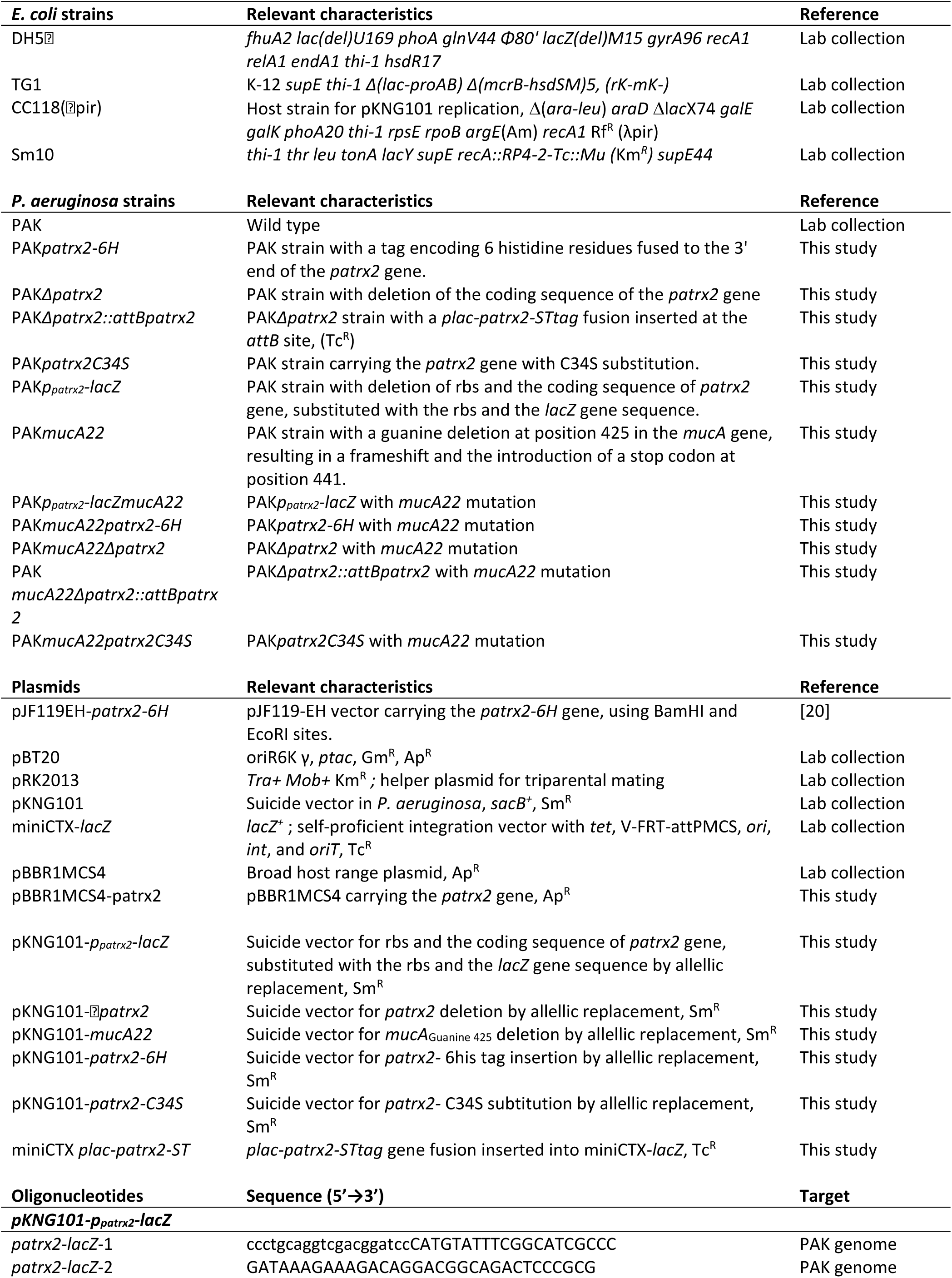
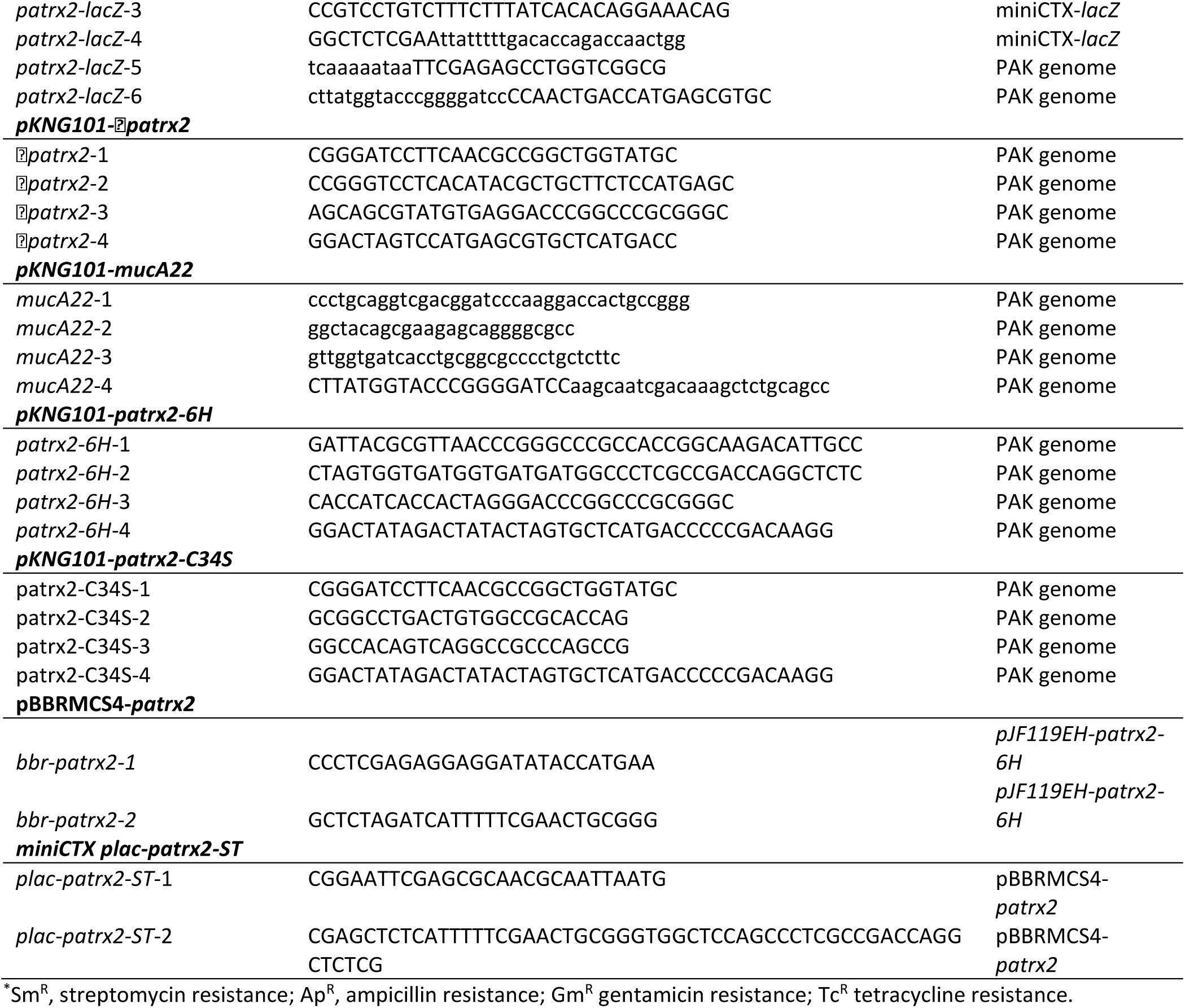
Bacterial strains, plasmids and oligonucleotides used in this study.

**Table S2:**
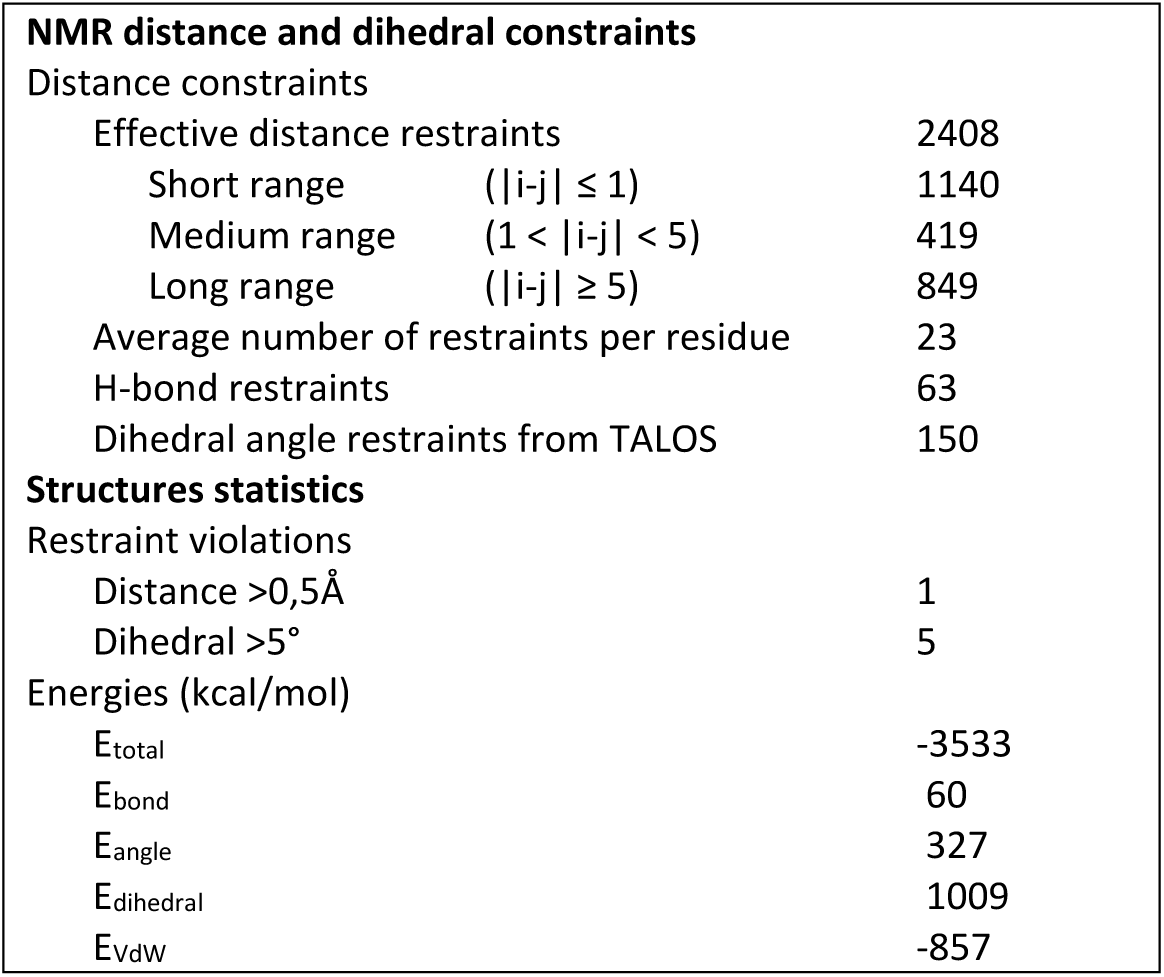
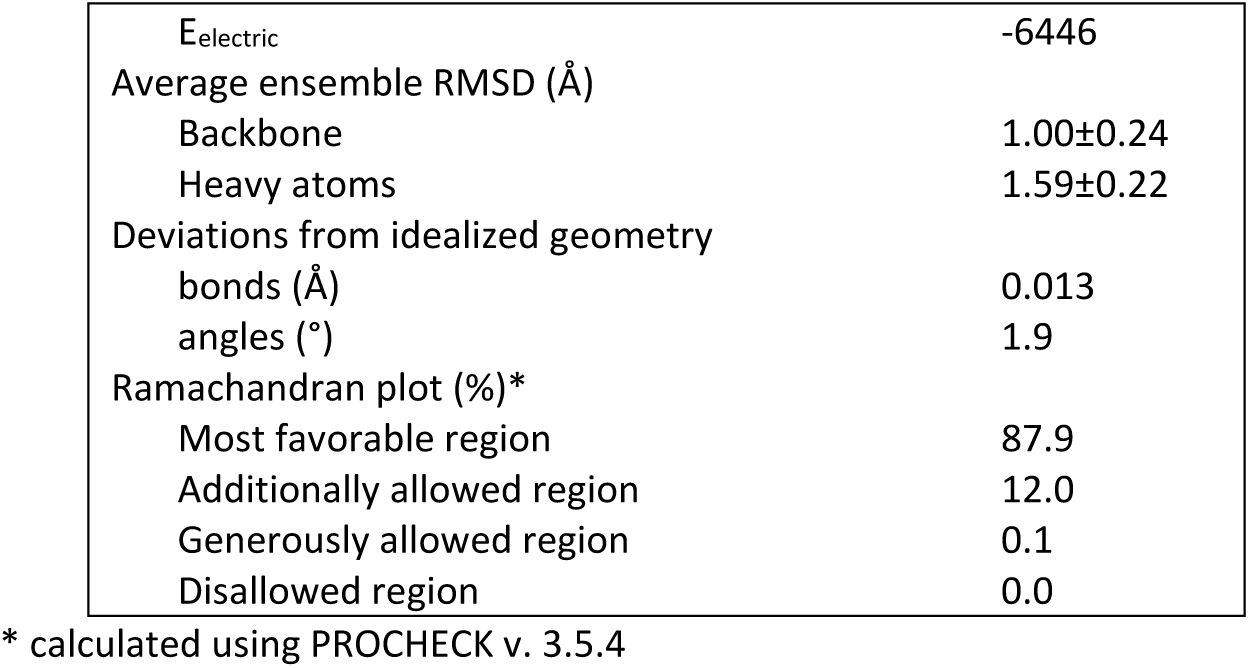
NMR and refinement statistics for Patrx2 structures. . Structural statistics and restraint violations of the 20 selected structures representative of Patrx2 in solution at pH 7.4 and 300K.

| NMR distance and dihedral constraints |  |  |
| --- | --- | --- |
| Distance constraints |  |  |
| Effective distance restraints |  | 2408 |
| Short range ($\vert i-j\vert \leq 1$) | | 1140 |
| Medium range ($1 < \vert i-j\vert < 5$) | | 419 |
| Long range ($\vert i-j\vert \geq 5$) | | 849 |
| Average number of restraints per residue |  | 23 |
| H-bond restraints |  | 63 |
| Dihedral angle restraints from TALOS |  | 150 |
| Structures statistics |  |  |
| Restraint violations |  |  |
| Distance >0,5Å |  | 1 |
| Dihedral >5° |  | 5 |
| Energies (kcal/mol) |  |  |
| E<sub>total</sub> |  | -3533 |
| E<sub>bond</sub> |  | 60 |
| E<sub>angle</sub> |  | 327 |
| E<sub>dihedral</sub> |  | 1009 |
| E<sub>vdw</sub> |  | -857 |

|  |  |
| --- | --- |
| E<sub>electric</sub> | -6446 |
| Average ensemble RMSD (Å) |  |
| Backbone | 1.00±0.24 |
| Heavy atoms | 1.59±0.22 |
| Deviations from idealized geometry |  |
| bonds (Å) | 0.013 |
| angles (°) | 1.9 |
| Ramachandran plot (%)* |  |
| Most favorable region | 87.9 |
| Additionally allowed region | 12.0 |
| Generously allowed region | 0.1 |
| Disallowed region | 0.0 |

\* calculated using PROCHECK v. 3.5.4

## Notes

### Competing Interest Statement

The authors have declared no competing interest.

